# Targeted degradation of the HPV oncoprotein E6 reduces tumor burden in cervical cancer

**DOI:** 10.1101/2024.10.17.618959

**Authors:** Tracess B Smalley, Angelo Nicolaci, Kim C. Tran, Jameela Lokhandwala, Nina Obertopp, Jenet K Matlack, Robert E Miner, Michael N. Teng, Shari Pilon-Thomas, Jennifer M Binning

**Author notes:** Correspondence should be addressed to Jennifer M. Binning.

## Abstract

Human Papilloma Virus (HPV)-related cancers are a global health burden, yet there are no targeted therapies available for chronically infected patients. The HPV protein E6 is essential for HPV-mediated tumorigenesis and immune evasion, making it an attractive target for antiviral drug development. In this study, we developed an E6-targeting Proteolysis Targeting Chimera (PROTAC) that inhibits the growth of HPV(+) tumors. To develop E6 antagonists, we generated a panel of nanobodies targeting E6 proteins derived from the oncogenic HPV16 subtype. The highest affinity E6 nanobody, A5, was fused to Von Hippel Lindau protein (VHL) to generate a PROTAC that degrades E6 (PROTAC^E6^). Mutational rescue experiments validated specific degradation via the CRL2^VHL^ E3 ligase. Intralesional administration of the PROTAC^E6^ using a clinically viable DNA vaccine reduced tumor burden in an immunocompetent mouse model of HPV(+) cancer. The inhibitory effect of the PROTAC^E6^ was abrogated by CD4^+^ and CD8^+^ T-cell depletion, indicating that the antitumor function of the PROTAC^E6^ relies in part on a host immune response. Overall, these results suggest that the targeted degradation of E6 inhibits its oncogenic function and stimulates a robust immune response against HPV(+) tumors, opening new opportunities for virus-specific therapies in the treatment of HPV-related cancers.

## Background

More than 80% of sexually active individuals will acquire human papillomavirus (HPV), making it the most prevalent sexually transmitted infection globally^1^. Persistent HPV infections cause up to six different cancers and are associated with nearly all cervical cancers and ∼25% of head and neck squamous cell carcinomas (HNSCCs)^2^. Despite the high efficacy of prophylactic HPV vaccines in preventing cervical cancer, HPV continues to contribute to 5% of the global cancer rates^3^. As there are currently no FDA-approved targeted therapies for the treatment of chronic HPV infections or HPV-associated malignancies, a large fraction of the global population remains at risk for HPV-related cancer^4^.

HPV-positive (HPV+) cancers depend on the expression of the viral E6 oncoprotein for tumor initiation, maintenance, and progression^5–9^. This dependency of the HPV lifecycle on E6 makes it an attractive target for therapeutic intervention^10–12^. E6 plays an important role in HPV pathogenesis and malignant transformation by interacting with multiple host proteins to transform homeostatic functions such as programmed cell death and cell cycle maintenance ^4,5,11^. It is well known that high-risk HPV E6 scaffolds p53 to the E3 ligase, E6-associated protein (E6AP), to ubiquitinate and degrade p53. The degradation of p53 blocks anti-proliferative and pro-apoptotic signaling, thereby altering the cellular environment to enhance viral genome amplification^4^. Additionally, E6 is involved in evading host immune responses, including downregulating the expression of interferon and interferon-stimulated genes to antagonize innate immune signaling pathways^13–15^. Given the multifunctional nature of E6 and its significance in HPV pathogenesis, this viral oncoprotein represents a key vulnerability that can be exploited in treating HPV-associated diseases.

Several approaches have been explored to therapeutically target E6. Targeting non-enzymatic proteins like E6 poses considerable challenges and often requires detailed information about protein-protein complexes and interfaces. Unlike enzymatic active sites, protein-protein interactions are often shallow with limited molecular features that can accommodate ligand binding. Nevertheless, considerable efforts have led to the development of a few small-molecule and antibody-based inhibitors, which have advanced our understanding of HPV biology and demonstrated the potential clinical benefits of targeting E6^16–19^. More recently, vaccine-based strategies have also been employed to target E6^20,21^. While these vaccines aim to induce cell-mediated immunity against E6 to eliminate infected cells, they do not directly inhibit the oncogenic functions of E6 and may be less effective in immunocompromised patients. Despite these advancements, no E6 antagonists have been approved for clinical use, underscoring the need for alternative strategies to target this oncoprotein.

Peptide PROTACs, or bispecific peptide-based degraders, are a promising technology for controlling endogenous protein expression through targeted ubiquitination^22^. Systems such as Biological Protein-Targeting Chimeras (bioPROTACs), Affinity-directed protein missile system (AdPROM), and Antibody RING mediated destruction (ARMeD), are engineered fusion proteins that consist of an E3 ligase or E3 ligase recognition protein and a target binding domain, which causes the ubiquitination and subsequent degradation of a protein of interest (POI)^23–27^. An advantage of peptide PROTACs is that they can engage the surface of a POI and are not limited to functionally relevant interaction sites. Therefore, peptide PROTACs offer a therapeutic approach for targeting proteins that are challenging to address with conventional small molecules, particularly multifunctional targets, as they enable the simultaneous neutralization of all functions through degradation.

Given the success of bioPROTACs and the challenges associated with targeting E6, we investigated whether E6 could be targeted for degradation using bioPROTAC technology. We developed a panel of high-affinity nanobodies against E6 and extensively characterized them for their ability to inhibit various oncogenic functions of E6. The highest affinity E6-binder was then developed into a bioPROTAC that promoted the degradation of E6. We found that the E6 bioPROTAC stimulated immune responses through the induction of interferon-stimulated genes (ISGs), and ultimately led to anti-tumorigenic effects in a pre-clinical setting. These data demonstrate the potential of bioPROTACs in targeting clinically relevant viral oncoproteins and underscore their broader implications for degrading other challenging therapeutic targets.

## Results

### Identification and characterization of nanobodies that bind HPV16 E6

HPV E6 is a relatively small protein composed of two zinc-binding domains, one E6AP-binding domain and one PDZ-binding domain^28^. Therefore, to isolate nanobodies against HPV16 E6, we purified a mutant form of E6 with enhanced expression and solubility^29^. Nanobodies were isolated from a yeast surface-displayed library of >1×10^8^ synthetic nanobody sequences (Fig 1A,B)^30^. Multiple rounds of selections, first by magnetic-activated cell sorting (MACS) and then by fluorescence-activated cell sorting (FACS), were carried out with biotinylated E6 (Fig. 1A,B). Four rounds of bio-panning yielded 11 unique nanobodies against HPV16 E6 (Fig 1A,B, Supp. Fig 1A). We performed an MBP pulldown assay to ensure the nanobodies bound to cellular E6. Multiple nanobodies pull down HPV16 E6 when co-expressed in *E. coli*, demonstrating their ability to engage E6 proteins in an intracellular environment (Fig. 1C). The binding affinity for our E6-targeting nanobodies was determined by microscale thermophoresis and revealed K_d_ values ranging from high picomolar to mid nanomolar (0.61 ± 0.7 – 458 ± 80 nM) (Fig 1D,E). Using differential scanning fluorimetry, we further determined that our nanobodies are highly thermostable, with melting temperatures greater than 46°C (Supp. Fig 1B).

**Figure 1.**
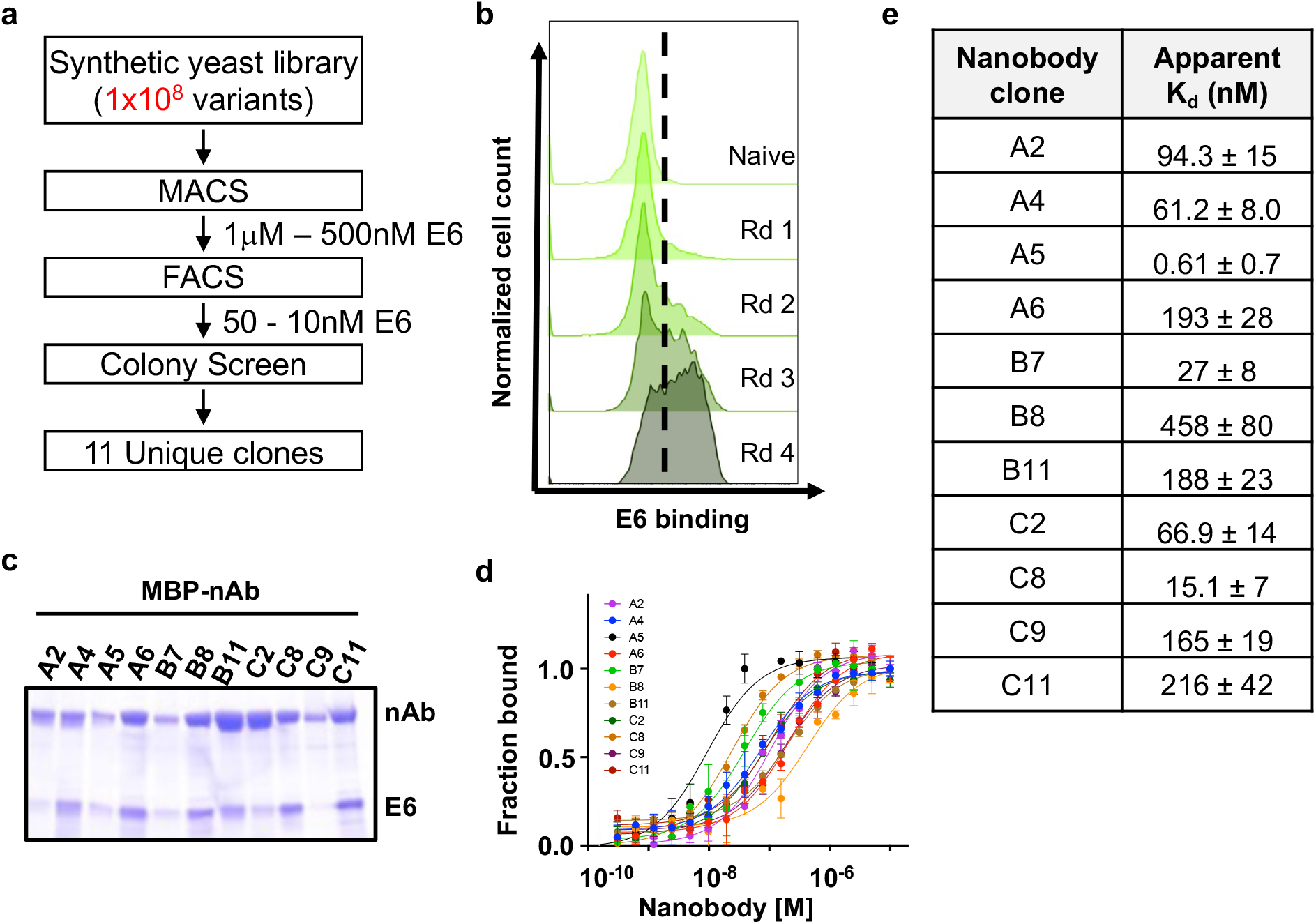
Generation and characterization of nanobodies to HPV16 E6. E6-specific nanobodies were discovered through **(a)** rounds of selection from a synthetic yeast surface display library, starting with enriching through magnetic cell sorting (MACS), fluorescent cell sorting (FACS), colony screen, and sequencing. **(b)** Flow cytometry histogram plots of yeast stained with fluorescently labeled E6 protein (50nM) following each round (Rd) of selection indicating enrichment in E6 binding. **(c)** Coomassie stained gels for MBP-E6 nanobodies fused co-expressed with His-E6 and pulled down with amylose resin. **(d)** Microscale thermophoresis (MST) binding curves for nanobodies interacting with the E6-sfGFP probe. Each point represents means of triplicates; error bars indicate standard deviation (SD). **(e)** Averaged binding affinities (K_D_) for all nanobodies as determined in **(d)**.

### Identification of nanobodies that inhibit HPV16 E6

We functionally categorized E6-binding nanobodies based on their ability to disrupt known E6 functions in HPV(+) CaSki cells. As E6 promotes uncontrolled cellular proliferation and the evasion of programmed cell death, two hallmarks of cancer, we initially assessed the ability of our nanobodies to inhibit colony formation and rescue apoptosis in the presence of E6^19^. In a colony formation assay, nanobodies A2 and C11 significantly reduced colony formation of CaSki cells (Fig 2A, Supp. Fig 2). To measure changes in apoptosis, we employed an Annexin V and propidium iodide (PI) assay (Supp. Fig 3A-D). A2 significantly restored apoptosis whereas C11 did not (Fig 2B,D Supp. Fig 3A). Importantly, the A2 nanobody did not induce apoptosis in HPV(-) C33A cells, indicating that the inhibitory properties of this nanobody are specific for HPV16 E6-expressing cells (Fig 2B,D, Supp. Fig 3B,D).

**Figure 2.**
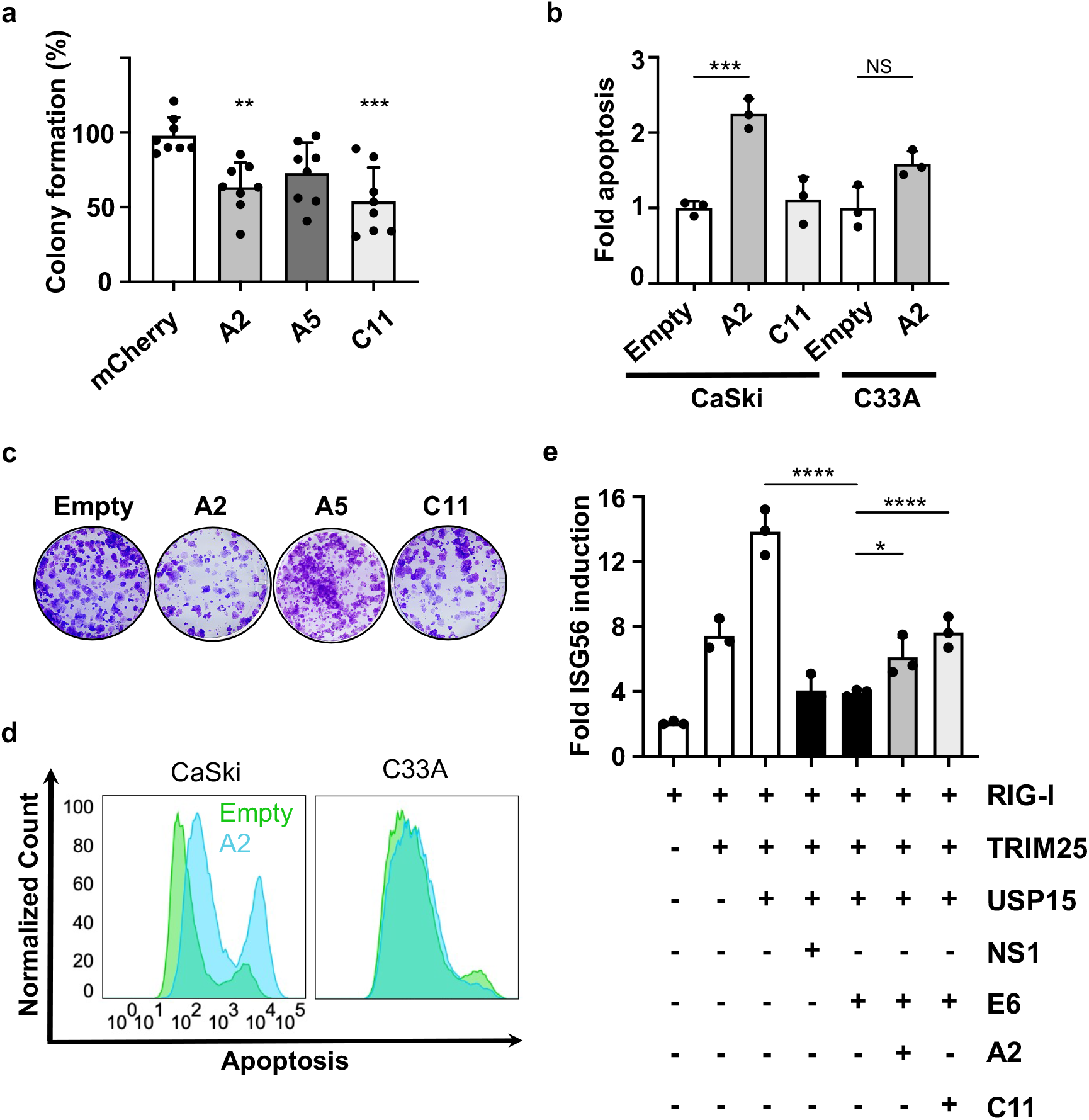
Nanobodies reduce colony-forming ability and induce apoptosis in HPV(+) cells. **(a)** Clonogenic assay. Colonies were stained with crystal violet and counted by Image J software. Colony-forming units were normalized to the mCherry control well in each plate and statistically analyzed using a one-way ANOVA (N=8). **(b)** Annexin V-APC/PI analysis of apoptosis compared to Empty control. Apoptosis was analyzed using FlowJo and analyzed with GraphPad using a one-way ANOVA (N=3). Error bars represent ± SD. **(c)** Colony forming assay images. **(d)** Annexin-V-APC/PI histograms for CaSki and C33A cell lines transfected with Empty control and A2 nanobody. **(e)** Dual Luciferase activity for ISG56 transcription. Luciferase activities shown are the means ± SD of triplicate samples for one of three representative experiments (N=3). (* p < 0.0332, ** p < 0.0021, *** p < 0.0002, **** p < 0.0001).

We next wanted to investigate how the E6-binding nanobodies influence host immune responses. We expected inhibitory nanobodies to activate innate immune signaling pathways and promote an antiviral immune state, given that E6 contributes to viral immune evasion^13,14,31,32^. We initially confirmed that HPV16 E6 antagonized innate immune signaling through the RIG-I pathway resulting in decreased transcription via the interferon-stimulated gene 56 (ISG56) promoter. We selected ISG56 as its transcription is strongly induced by interferons and viral infection, and it has been reported to inhibit HPV replication^33^. Importantly, E6 expression significantly reduced ISG56 transcription to levels comparable to RSV NS1, a well-established RIG-I antagonist^34^ (Fig 2E, Supp. Fig 4A). Expression of A2 and C11 counteracted the immune antagonistic function of HPV16 E6 by restoring ISG56 activation by ∼50% or 100%, respectively. Further, C11 restored the induction of ISG56 in a dose-dependent manner (Supp. Fig 4B), highlighting the inhibitory effects of this nanobody. Together, these data establish that we have isolated inhibitory nanobodies that disrupt multiple E6 functions and that each nanobody has a distinct mechanism of inhibition.

### Targeted degradation of HPV E6 by a VHL-based PROTAC

As an additional strategy for inhibiting E6, we designed a bioPROTAC that targets E6 for proteolytic degradation (PROTAC^E6^). We hypothesized that an E6 degrader could disrupt several E6 functions more effectively than nanobodies alone, since the latter may only target a small binding footprint within the overall E6 structure. The PROTAC was generated by fusing our highest affinity E6-binding nanobody, A5, to the Von Hippel Lindau protein (VHL). This format is designed to mediate the degradation of E6 via recruitment to the Cullin2 (CUL2) Ring E3 ligase complex (CRL2) bound by VHL (CRL2^VHL^)^35–37^ (Fig 3A). To assess the effects of PROTAC^E6^, we expressed E6-GFP alone and in combination with VHL, PROTAC^E6^, or a PROTAC^E6^ mutant that abolishes CUL2 binding (mutPROTAC^E6^)^38^. We found that PROTAC^E6^ expression resulted in extensive degradation of E6-GFP (based on GFP detection), whereas the VHL control did not. CRL2 recruitment was critical for E6 degradation as the expression of the mutPROTAC^E6^ prevented E6 degradation (Fig 3B). The functionality of the PROTAC^E6^ was further verified by assessing its effect on the proliferation of E6-dependent CaSki cells. The PROTAC^E6^ inhibited cellular proliferation in a dose-dependent manner (Fig 3C) and had a more significant reduction of cellular proliferation when compared to the A2 and C11 nanobodies (Fig 3D). These data demonstrate that PROTAC^E6^ can efficiently target E6 for degradation in a CRL2-dependent manner.

**Figure 3.**
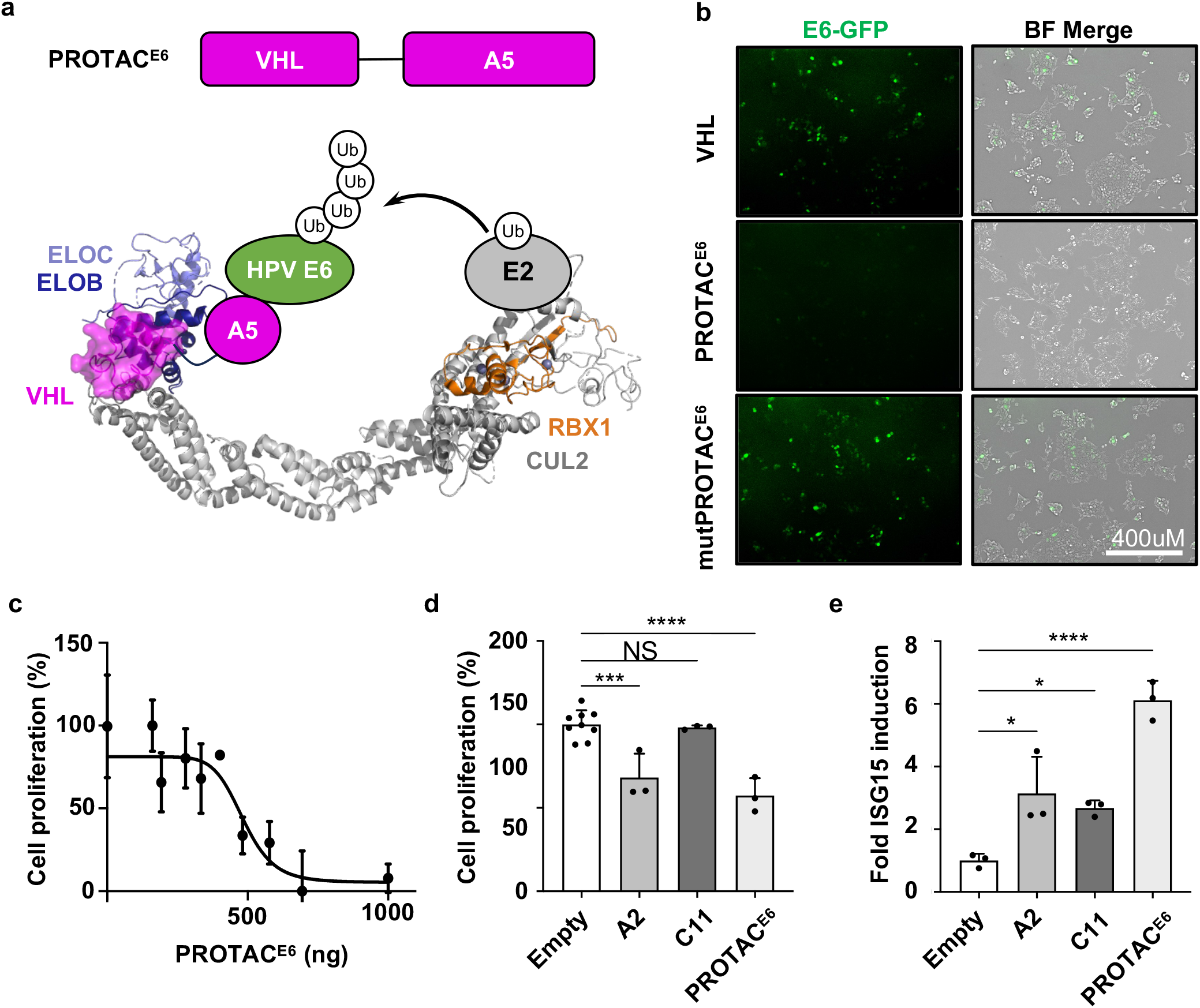
BioPROTAC^E6^ degrades E6, disrupts HPV(+) proliferation and induces ISG15 mRNA. **(a)** Schematic representing PROTAC^E6^ interacting with CUL2. **(b)** Confocal microscopic images of HEK293T cells transfected with E6-GFP alone, or in combination with VHL, PROTAC^E6^(VHL-A5), or mutPROTAC^E6^ (VHL^TLK157-159AAA^-A5) and imaged after 48 hours (N=3). **(c)** The dose-dependent titration of PROTAC^E6^ was assessed for cellular proliferation in CaSki cells 48 hours post-transfection, using WST-1 reagent in a 96-well plate (N=4, error bars ± SD). A curve was fitted using Prism GraphPad. **(d)** CaSki cells transfected with Empty control, A2, C11, and PROTAC^E6^ were assessed 48 hours later for cellular proliferation using WST. **(e)** Induction of ISG15 mRNA using RT-qPCR (N=3). A one-way ANOVA was performed, and error bars represent ± SD. (* p < 0.0332, ** p < 0.0021, *** p < 0.0002, **** p < 0.0001).

### Nanobodies and PROTAC^**E6**^ **restore innate immune signals in HPV(+) cells**

To investigate the effect PROTAC^E6^ has on the innate immune response, we measured the transcription of ISG15, another interferon-stimulated gene that plays a role in mediating an anti-viral response. We found that A2, C11, and PROTAC^E6^ each increased the mRNA levels ISG15 in CaSki cells (Fig 3E). The most substantial effect was observed using the PROTAC^E6^, which increased ISG15 levels by > 6-fold compared to untreated cells (Fig 3E). Additionally, PROTAC^E6^ restored the transcription of IFNβ in CaSki cells (Supp. Fig 5). These data demonstrate that the nanobodies can inhibit E6 and rescue innate signaling, and that the PROTAC^E6^ was a more efficient E6 antagonist than either nanobody alone.

### In vivo depletion of HPV16 E6 inhibits tumor growth

Based on the in vitro effectiveness of the PROTAC, we sought to assess the therapeutic potential of the PROTAC^E6^ administration in a mouse model of HPV(+) cancer. For this approach, we developed a PROTAC-encoding DNA vaccine for intralesional delivery. We selected the pAC vector based on its efficacy as a DNA vaccine and previous use in clinical trials^39–41^. We initially confirmed that PROTAC^E6^ treatment inhibits cell growth in TC-1 cells, a mouse cell line that stably expresses HPV16 E6 and E7 (Fig 4A). The TC-1 tumor cells were subcutaneously injected into the ventral position of C57BL/6 mice. Once tumors were established, the PROTAC^E6^ was administered by intralesional injections on days 7 and 14 (Fig 4B). By the study endpoint, PROTAC-mediated degradation of E6 decreased the tumor volume by over 50% (Fig 4C).

**Figure 4.**
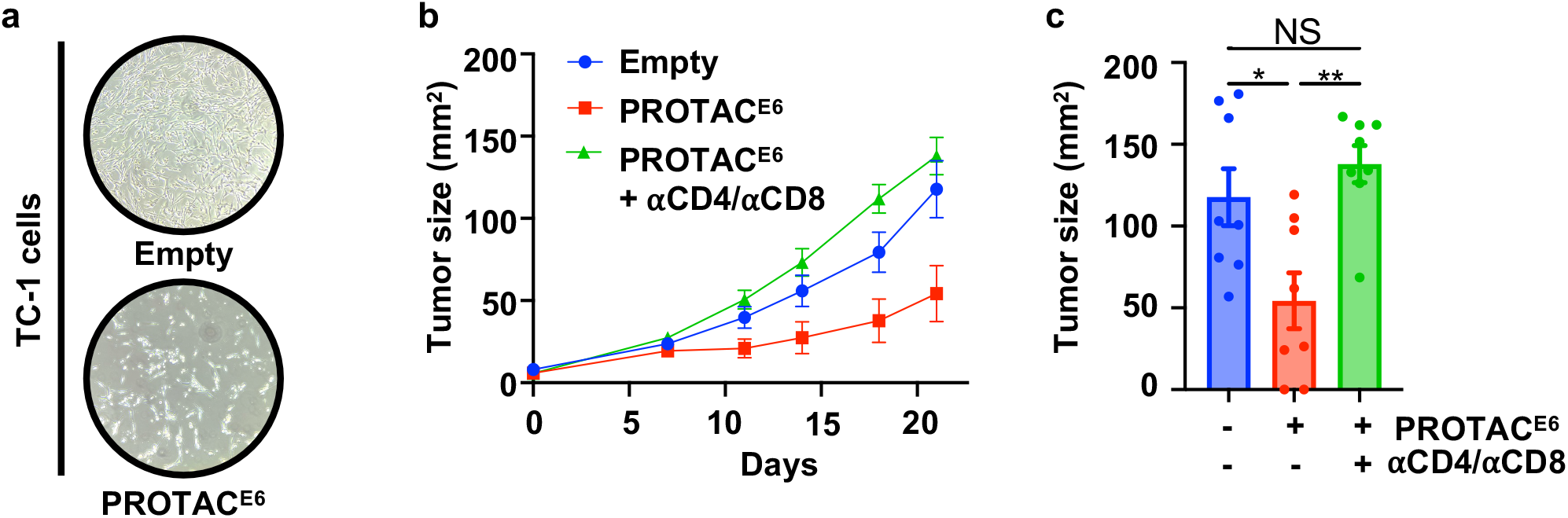
PROTAC^E6^ mediated the reduction in TC-1 tumor burden. **(a)** Brightfield microscopic images of TC-1 cells transfected with Empty and PROTAC^E6^ and imaged 48 hours later. **(b)** Tumor growth curves after treatment on days 7 and 14 with Empty (N=8) and PROTAC^E6^ (N=16) and a subgroup on PROTAC^E6^ treated with αCD4 and αCD8 (N=8). **(c)** Endpoint tumor size was graphed and analyzed using a one-way ANOVA and error bars represent ± SEM. (* p < 0.0332, ** p < 0.0021, *** p < 0.0002, **** p < 0.0001).

Given that E6 inhibition increases interferon stimulation coupled with the degradation-based mechanism of PROTACs, we hypothesized that the E6 PROTAC could also contribute to immune clearance by promoting the presentation of foreign peptides to immune cells in the tumor microenvironment. To investigate this possibility, we tested whether the depletion of T cells would influence the antitumor effect of the PROTAC. We found that the PROTAC^E6^ indeed relied on the host immune response, as depletion of CD4^+^ and CD8^+^ cells abrogated PROTAC^E6^-mediated inhibition of tumor growth (Fig 4B,C). Importantly, no weight loss or adverse effects were observed in PROTAC^E6^ treated mice (Supp. Fig 6). These results establish that degrading E6 is a viable therapeutic modality to reduce tumor burden and stimulate an anti-tumor immune response.

## Discussion

Our study demonstrates the effectiveness of E6 degradation in inhibiting the growth of HPV(+) tumors. Thus far, the therapeutic potential of PROTACs has been greatly limited due to their inability to cross the cell membrane. This limitation holds true for both protein-based PROTACs and bulky, bifunctional chemical PROTACs with suboptimal pharmacological properties. Here, we overcome this hurdle using a DNA vaccine method that is uniquely suited for targeting visible lesions. Although this method is viable for nearly all HPV(+) tumors due to their localization patterns, our data shows how the potential of protein-based PROTACs can be unlocked in certain contexts using the appropriate delivery system.

The preliminary success of this approach warrants further optimization of both the vaccine vector and composition. The HPV proteins E6 and E7 are both critical for HPV-mediated tumorigenesis and immune evasion. Therefore, developing PROTACs for more than one HPV oncoprotein (i.e. E6 and E7) may further improve tumor-growth inhibition by antagonizing the function of multiple proteins. Alternatively, the incorporation of chemokines or cytokines that enhance immune infiltration may also increase efficacy given the demonstrated requirement for T cells for PROTAC antitumor function in vivo. Encapsulation of the DNA vaccine in nanoparticles or other biomaterials may also facilitate uptake or delivery. Regardless, the tumor inhibition achieved through the current method provides a strong foundation for the development of future therapeutic candidates.

Our study also identified that nanobodies A2 and C11, restrict E6 oncogenic functions and have distinct inhibitory properties. Specifically, A2 induces apoptosis in HPV(+) cells, whereas C11 does not. Conversely, C11 restores innate immune signaling, while A2 does not. Although the binding epitopes of A2 and C11 are not yet identified, they likely engage unique regions of E6 based on their ability to disrupt distinct E6 functions. This observation aligns with known separation-of-function mutations in E6, such as mutations that disrupt interactions with either E6AP, P53, or NHERF1^42,43^. Importantly, biological tools that bias E6 functions can be leveraged to help identify which E6 functions are essential at different stages of viral infection and cancer progression.

Collectively, this study generated a diverse panel of nanobodies against HPV16 E6 and demonstrated that E6-targeting PROTACs can activate host immune responses and inhibit tumor growth. These findings establish a precedent for developing E6-degrading therapeutics and show the effectiveness of delivering bioPROTACs via intralesional injection. More broadly, this research highlights the therapeutic potential of targeted protein degradation for addressing clinically relevant, yet challenging, targets.

## Methods

### Plasmids

All constructs were generated by standard PCR and restriction-based cloning methods unless otherwise noted. The HPV16E6^4C/4S^ was codon optimized for bacterial expression (Integrated DNA Technologies) and cloned into a modified pET15b vector containing an N-terminal MBP-tag, His-tag, and tobacco etch virus (TEV) cleavage site with and without a sfGFP fusion (HPV16 E6-sfGFP)^44^. The E6 nanobodies were cloned into the pET26b (+) periplasmic expression vector as previously described^30^. For mammalian expression, A2, A5, C11, VHL-A5 (PROTAC^E6^), and mutPROTAC^E6^ were cloned into pcDNA3.1-3xHA. For in vivo intralesional injections (IL), PROTAC^E6^ was cloned into the pAc vector containing a C-terminal 3xHA-tag. All constructs were sequence validated.

### Protein expression and purification

All proteins were expressed in *E. coli*. Plasmids were transformed into *E. coli* BL21(DE3) (Invitrogen) cells and grown at 37°C to an optical density of 0.6–0.8 and induced with 0.5 mM isopropyl β-D-1-thiogalactopyranoside (IPTG) overnight at 18°C. For HPV16 E6 and HPV16E6-GFP, cells were harvested and subsequently lysed in buffer containing 50 mM phosphate pH 6.5, 150 mM NaCl,1 mM dithiothreitol [DTT], 20 mM imidazole and 10% glycerol, followed by centrifugation (40,000× g, 30 min, 4°C) to clarify the lysate. The sample was purified by nickel affinity chromatography and cation (SP) exchange columns. After cation exchange, the MBP and 6xHis tags were proteolytically removed by TEV treatment. The sample was purified by a GE Superdex 75™ increase 10/300 size exclusion column (SEC) into a buffer containing 50 mM phosphate pH 6.5, 150 mM NaCl, 1 mM TCEP.

### Periplasmic expression and purification of Nbs

Nanobodies were cloned into the periplasmic expression vector pET26b (+) and purified as described previously^30^. Briefly, nanobodies were grown in *E. coli* and lysed after overnight induction at 18°C. Nanobody-containing supernatant was incubated with 2.5 mL of Ni-NTA Resin (Thermofisher) and 0.01% NaN_3_ overnight at 4°C. The Ni-NTA resin was then washed with a high-salt buffer (20 mM Tris, pH 7.5, 500 mM NaCl, 20 mM Imidazole), followed by a wash with nickel wash buffer (20 mM Tris, pH 7.5, 150 mM NaCl, 20 mM imidazole). The nanobodies were eluted with nickel elution buffer (20 mM Tris, pH 7.5, 150 mM NaCl, 250 mM Imidazole). SEC was used as a final purification step and the nanobodies were stored in buffer containing 20mM HEPES pH 7.5, and 150 mM NaCl. The melting temperature for the nanobodies was measured by differential scanning fluorimetry using 5µM of protein and 5XSYPRO™ dye (ThermoFisher S6650).

### Bio-panning for HPV16 E6-binding Nbs using yeast surface display

We used a previously published yeast synthetic nanobody library with a diversity of 10^8^ variants to identify nanobodies against viral proteins^30^. Nanobodies were isolated using yeast display panning protocol described in detail^45^. Briefly, purified E6 was N-terminally biotinylated by incubation with a 5X molar excess of Sulfo-NHS-Biotin salt (Thermo) overnight at 4°C. Biotinylated protein was purified via SEC into a buffer containing 20 mM HEPES, pH 7.4, 150 mM NaCl, 1 mM TCEP and stored at −80 °C. The panning was accomplished in four rounds using two rounds of magnetic activated cell sorting (MACS) at 1µM and 500 nM and two rounds of fluorescent activated cell sorting (FACS) at 50 nM, and 10 nM of biotinylated E6 protein. Negative selection was performed before each round of selection by incubating yeast with Alexa Fluor 647-conjugated streptavidin (SA-647) and anti-CY5/647 microbeads (Miltenyi). MACS was performed by incubating biotinylated protein with streptavidin-coated microbeads (Miltenyi) at 4 °C for 30 min, subsequently flowed over a MACS LS separation column (Miltenyi) pre-equilibrated in selection buffer (20 mM HEPES pH 7.4, 150 mM NaCl, 5% Maltose) and analyzed using an Accuri Flow Cytometer (BD Accuri C6 Plus). Positive selections for FACS were achieved by incubating the yeast with biotinylated HPV16 E6 and staining with Alexa Fluor 647-conjugated streptavidin and Alexa Fluor 488-conjugated anti-HA mouse monoclonal antibody (Cell Signaling) and sorting using a FACS SH800S Cell sorter (Sony). The isolated yeast populations obtained after round 4 were subjected to colony screening and plasmid recovery using a Zymoprep yeast plasmid kit (Zymo Research). The plasmids were transformed into *E*. coli and 30 individual colonies were sequenced with primer pYAL F (Eurofins). Sequencing data analysis was performed using SnapGene software (Dotmatics).

### In-vitro pull down assays

MBP-fused nanobody constructs were co-transformed with His-E6 in *E. coli* cells (BL21 DE3), grown until OD 0.6, and induced overnight at 18°C for 18 hours. Cells were lysed with buffer (20 mM HEPES, pH 7.4, 150 mM NaCl, 0.5% IGEPAL CA-630, 1mM DTT) and incubated with amylose (New England Biolabs) resin for 20 minutes at room temperature. The resin was washed three times with buffer and collected for SDS PAGE analysis.

### Microscale Thermophoresis (MST) binding assay

The binding affinities of the E6 nanobodies were measured using microscale thermophoresis with a Monolith NT.115 machine (NanoTemper Technologies). For binding affinity determination, 50 nM of HPV16 E6-sfGFP in HBS-T (20 mM HEPES pH 7.5, 150mM NaCl, 1mM DTT, 10 mM MgCl_2_, 0.05 % (v/v) Tween 20 and 0.1% (w/v) BSA) was mixed in equal parts with serial-diluted E6 nanobodies in HBS-T producing final ligand concentrations ranging from 20 μM to 150 pM. The HPV16 E6-sfGFP/E6 nanobodies mixture was incubated for 10 min at an ambient temperature of 25°C prior to mounting them into standard capillaries (MO-K022, NanoTemper Technologies) and using the NanoBlue filter to excite the fluorophore to measure MST traces. Three independent experiments were performed at 50% LED power, medium MST power. The data was analyzed using the NTAnalysis software (NanoTemper Technologies) and figures were curve-fitted using GraphPad Prism.

### Cell culture and transfections

CaSki HPV(+) cervical cancer cells were cultured in RPMI 1640 (1X) supplemented with 10% FBS, 100 units/ mL of penicillin, 100mg/mL of streptomycin, 2mM L-glutamine, 1 mM sodium pyruvate, NEAA 0.1mM, 0.05mM 2-mercaptoethanol, 0.5mg/mL amphotericin B and 0.5mg/mL gentamycin. C33A HPV(-) cervical cancer cells and HEK293T were cultured in DMEM (1X) supplemented with 10% FBS, 100 units/ mL of penicillin, 100mg/mL of streptomycin, and 0.5mg/mL amphotericin B. CaSki (ATCC, CRM-CRL-1550), HEK293T (a generous gift from the Lau lab) and C33A (ATCC, HTB-31) were kept at passages less than 10, mycoplasma tested quarterly and maintained in an incubator at 37 °C and 5% CO_2_ atmosphere. Transient transfections were performed with poly (ethylenimine) (PEI 1mg/mL, 4:1 reagent to [DNA]), according to manufacturer’s protocols.

### Colony formation screen

CaSki cells were seeded into 6-well plates at 3×10^5^ cells per well and transfected 24 hours later. 48 hours after transfections, CaSki cells were trypsinized and redistributed into 6-well plates at 800 cells per well. Cells were allowed to grow for 2-3 weeks, and subsequent cells were fixed with 100% methanol, and stained with 0.5% (w/v) crystal violet. Colony forming images were processed using ImageJ Fiji expansion^46^.

### Apoptosis screen

CaSki cells were seeded into 6-well plates at 3×10^5^ cells per well and transfected 24 hours later. Cells were harvested after 48 hours, washed with PBS and stained with Annexin-V-APC and propidium iodide (500ng/10^6^ cells) according to manufacturer’s instructions. Late-stage apoptosis was analyzed within 1 hour of staining using a BD FACS Celesta.

### ISG56 Activation assay

HEK293T cells were plated into 24-well at 1.6 ×10^5^ cells per well and incubated overnight at 37°C. The next day, expression plasmids encoding constitutively active RIG-I (N-RIG-I, 4 ng), TRIM25 (100 ng), USP15 (62 ng), plus a firefly luciferase reporter under the control of the ISG56 promoter (pISG56-luc, 20 ng) were transfected into the cells using GeneJuice (EMD Millipore). The phRL-TK (5 ng) (Promega), expressing the Renilla luciferase gene under the control of the HSV-TK promoter, was used as a transfection control. Expression plasmids encoding HPV16 E6 (200 ng) or the RSV NS1(100 ng) control were included to inhibit RIG-I signaling; plasmids encoding the E6-targeting nanobodies (400 ng) were included with HPV16 E6 as indicated. Cells were harvested 24h post-transfection and subjected to Dual-Luciferase assay (Promega). Specifically, 100µl of Passive Lysis Buffer was added per well and allowed to incubate for 30 minutes. Then 30µl of the clarified lysate was transferred to a well of an opaque white 96-well plate (Greiner Bio One). The Luciferase Assay Reagent II (LARII) and Stop & Glo Reagent were dispensed 50uL per well and relative firefly and Renilla luciferase activity were measured using the BioTek Synergy MX plate reader. Samples were done in triplicate.

### Quantitative real-time PCR

IFNB and ISG15 expression was analyzed in CaSki cells using quantitative PCR. CaSki cells were seeded into 6-well plates at 3×10^5^ cells per well and transfected with 2.5 µg of empty, A2, C11 and PROTAC^E6^ constructs 24 hours later. Cells were stimulated with polyinosinic-polycytidylic acid sodium salt (poly (I:C)) for 6 hours prior to harvest and TRIzol was added for mRNA extraction. mRNA extraction was performed according to the manufacturer’s protocol (Invitrogen TRIzol Reagent, 15596026) and RNA quality was assessed using NanoDrop 2000c Spectrophotometer (ThermoFisher). Initially, reverse transcription was performed using cDNA reverse transcription (Applied Biosystems™ High-Capacity cDNA Reverse Transcription Kit, 4368814) and DNA quantification was analyzed using commercially available SYBR green (SYBR Green PCR Master Mix, 4344463). Primers are listed in primer table. Gene expression levels were normalized to cellular GAPDH. Fold induction of each target gene relative to Empty treated cells was calculated using the delta delta CT method (ΔΔC_T_). RT-qPCR assays were recorded on StepOnePlus Real-Time PCR System (Applied Biosystems).

### PROTAC reporter assay

HEK293T cells were seeded into 6 well plates at a cell density of 3×10^5^ cells per well and transfected with 1.25µg of the construct E6-GFP and co-transfected with 1.25µg of VHL, PROTAC^E6^ and mutPROTAC^E6^ as described above. After 48 hours post-transfection live cell imaging was performed using an EVOS FL Fluorescent Microscope (Invitrogen) to monitor the expression of E6-GFP.

### Cell proliferation assays

CaSki cells were seeded into 96-well plate at 1×10^3^ cells per well with 200µL of media. For comparison assays, cells were treated with 700ng of Empty, A2, C11 and PROTAC^E6^. For the dose-dependent assay, cells were treated with different concentrations of PROTAC^E6^ (0, 161, 193, 279, 334, 401, 482, 578, 694, and 1000 ng, N=3), but all cells received 1000 ng total of DNA with the addition of Empty vector. After 48 hrs of treatment, the cell proliferation was analyzed using Water-Soluble Tetrazolium (WST-1, Roche, 5015944001) at wavelengths 450 and 600 nm. The absorbance of 450 nm was corrected by subtracting the absorbance 600 nm. The plates were read on a BioTek SynergyHT microplate reader (Agilent Technologies). Cell proliferation was graphed in GraphPad 9.0 and normalized to transfection efficiency.

### PROTAC inhibition of E6 tumor progression in mice

Female C57BL/6 mice (6–8 weeks old) were purchased from Charles River and housed at the Animal Research Facility of the H. Lee Moffitt Cancer Center and Research Institute (Tampa, FL). TC-1 tumor cells were injected subcutaneously and ventrally in C57BL/6 mice. Seven days after TC-1-bearing mice exhibited palpable tumors (approximately 25 mm^2^), mice received two doses of IL injection of Empty or PROTAC^E6^ cloned into a previously published pAc vector at weekly intervals. The Empty and PROTAC^E6^ vectors were prepared with *Invivo*-jetPEI transfection reagent (PolyPlus Transfections) by mixing 20μg of Empty (control, N=8), and PROTAC^E6^ (N=16), and injected intratumorally on days 7 and 14. A CD4 and CD8 depletion subgroup of the PROTAC^E6^ vector (N=8) was treated with α-CD4 and α-CD8 at a concentration of 300µg/ 200µL/mouse/i.p three days prior to the first injection, and subsequently twice per week during the duration of the experiment. Tumor growth was measured two times per week and harvested on day 21.

## Statistical analysis

Statistical comparisons were carried out using GraphPad Prism 9.0. MST and dose-dependent titration binding curves were fitted using GraphPad Prism by performing ([Agonist] vs response (three parameters) and [Inhibitor] vs. response –variable slope (four parameters), respectively). One-way ANOVA was utilized for colony formation, apoptosis, ISG56 induction, qPCR results, and analysis of mouse tumor burden.

---

**Supplemental figure 1. Biophysical characterization of full nanobody panel. (a)** Gating for yeast surface display selections. **(b)** Tabulated averaged melting temperatures for all nanobody candidates are represented in a table ± SD.

**Supplemental figure 2. Quantified colony formation screen. (a)** Bar graph of full colony formation screen for all 11 nanobodies, analyzed using a one-way ANOVA (N=8, error bars ± SD). (* p < 0.0332, ** p < 0.0021, *** p < 0.0002, **** p < 0.0001).

**Supplemental figure 3. Apoptosis screen for full panel of nanobodies. (a)** Apoptosis screen for all 11 nanobodies in **(a)** CaSki and **(b)** C33A was analyzed using a one-way ANOVA (N=3, error bars ± SD). **(b)** Significant nanobodies from the CaSki screen were further screened in C33A HPV(-) cervical cancer cells (N=3, error bars ± SD) and analyzed with a one-way ANOVA. **(c)** Flow cytometry gating schema for the apoptosis screen. **(d)** Representative dot plots for CaSki and C33A cell lines in apoptosis screen with Empty control and A2 nanobody. (* p < 0.0332, ** p < 0.0021, *** p < 0.0002, **** p < 0.0001).

**Supplemental figure 4. ISG56 screen and dose-dependent relationship of nanobody C11**. HEK293T cells were transfected with expression plasmids encoding N-RIG-I, TRIM25, USP15, pISG56-luc, and HPV16 E6 without and with **(a)** the full range of nanobodies or **(b)** increasing concentrations of the C11 E6-targeting nanobodies as indicated. Luciferase activities were normalized by co-transfection with phRL-TK. Cell lysates were collected after 24 h and luciferase activities were determined by Dual Luciferase Assay (Promega). Shown are the means ± SD of triplicate samples for one of three representative experiments (N=3).

**Supplemental figure 5. PROTAC**^**E6**^ **induction of IFNB mRNA levels**. The induction of IFNB mRNA was measured using RT-qPCR (N=3). A one-way ANOVA was performed, and error bars represent ± SD. (* p < 0.0332, ** p < 0.0021, *** p < 0.0002, **** p < 0.0001).

**Supplemental figure 6. Mouse body weight**. Mouse body weight was recorded throughout treatment and plotted (N=5).

## Data Availability Statement

All data needed to reproduce the results are available within the manuscript supplementary information and source data provided with the paper. Other relevant data is available from the corresponding authors. Source data is provided with this paper.

## Ethics Statement

The authors declare no competing interests.

## Author Contributions

T.B.S performed all differential scanning fluorimetry, cell-based assays, mouse studies, contributed to the experimental design, and wrote the manuscript.

A.N. performed E6 nanobody selections, protein purifications, and microscale thermophoresis binding assays, contributed to experimental design and wrote and edited the manuscript.

N.O. assisted with mouse experiments, and flow staining and edited the manuscript.

J.K.M. assisted with mouse experiments, and flow staining and wrote the manuscript.

R.E.M. developed the mutPROTAC^E6^ construct and edited the manuscript.

J.L. assisted with data collection and wrote and edited the manuscript.

K.C.T. developed and performed ISG56 dual luciferase assay and wrote the manuscript.

M.N.T. provided resources, reviewed and edited the manuscript. S.PT supervised, provided resources, and reviewed the manuscript.

J.M.B. conceptualized, supervised, provided resources, reviewed, and edited the manuscript. All authors read and approved the final manuscript.

## Acknowledgements

The authors would like to thank Luca lab members for their suggestions and technical assistance, Dr. Andy Kruse for the nanobody library, and Dr. Eric Lau’s lab for the HEK293T cell line. The authors also acknowledge the Analytical Microscopy Core and the Chemical Biology Core Facilities at the H. Lee Moffitt Cancer Center & Research Institute, which is supported in part by a National Cancer Institute (NCI) support grant (P30-CA076292).

## Supplemental Material

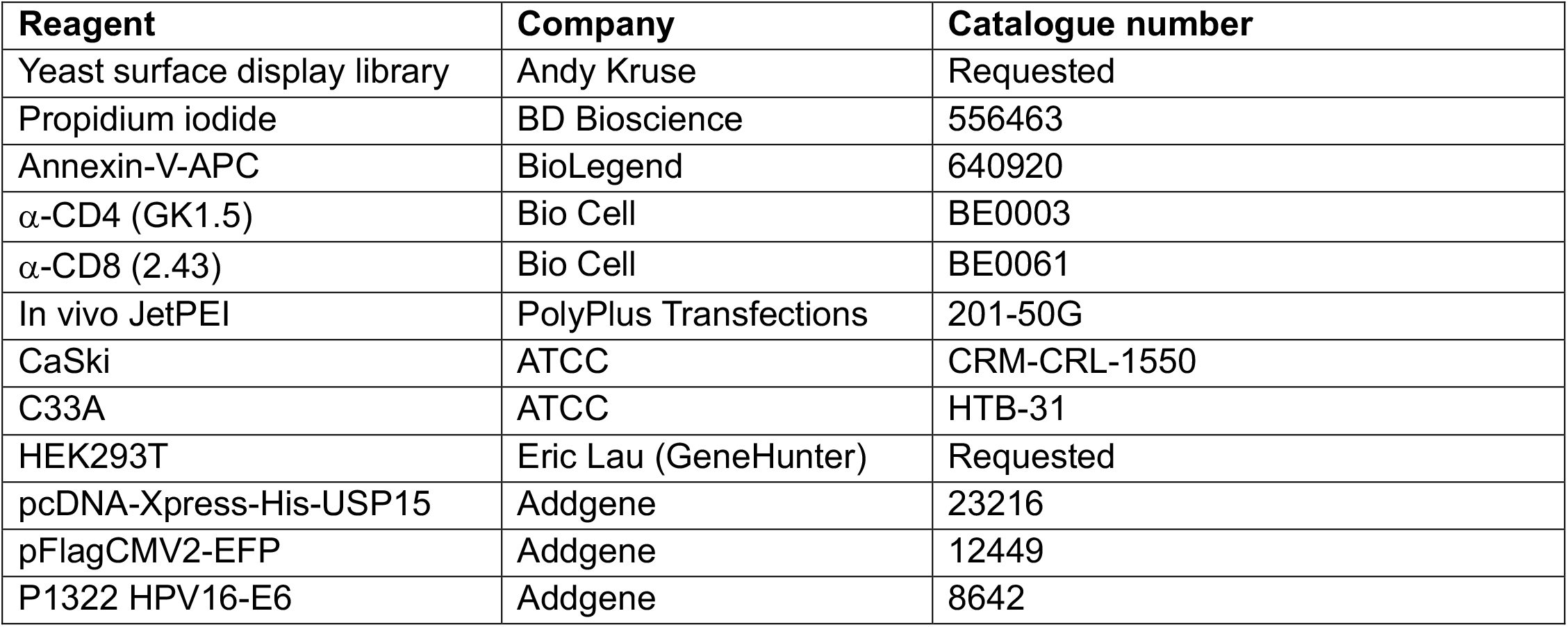

| Reagent | Company | Catalogue number |
| --- | --- | --- |
| Yeast surface display library | Andy Kruse | Requested |
| Propidium iodide | BD Bioscience | 556463 |
| Annexin-V-APC | BioLegend | 640920 |
| α-CD4 (GK1.5) | Bio Cell | BE0003 |
| α-CD8 (2.43) | Bio Cell | BE0061 |
| In vivo JetPEI | PolyPlus Transfections | 201-50G |
| CaSki | ATCC | CRM-CRL-1550 |
| C33A | ATCC | HTB-31 |
| HEK293T | Eric Lau (GeneHunter) | Requested |
| pcDNA-Xpress-His-USP15 | Addgene | 23216 |
| pFlagCMV2-EFP | Addgene | 12449 |
| P1322 HPV16-E6 | Addgene | 8642 |

## Table of primers

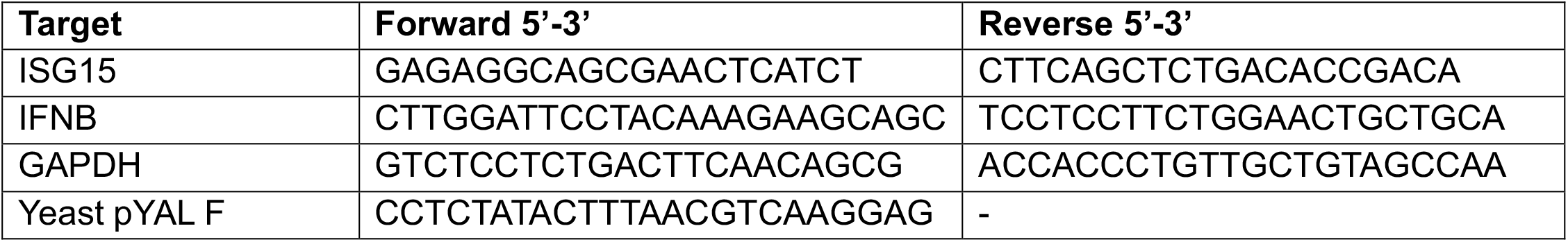

