## Supplemental_files for "Targeted degradation of the HPV oncoprotein E6 reduces tumor burden in cervical cancer"

Supplemental figure 1 . Biophysical characterization of full nanobody panel

**a**

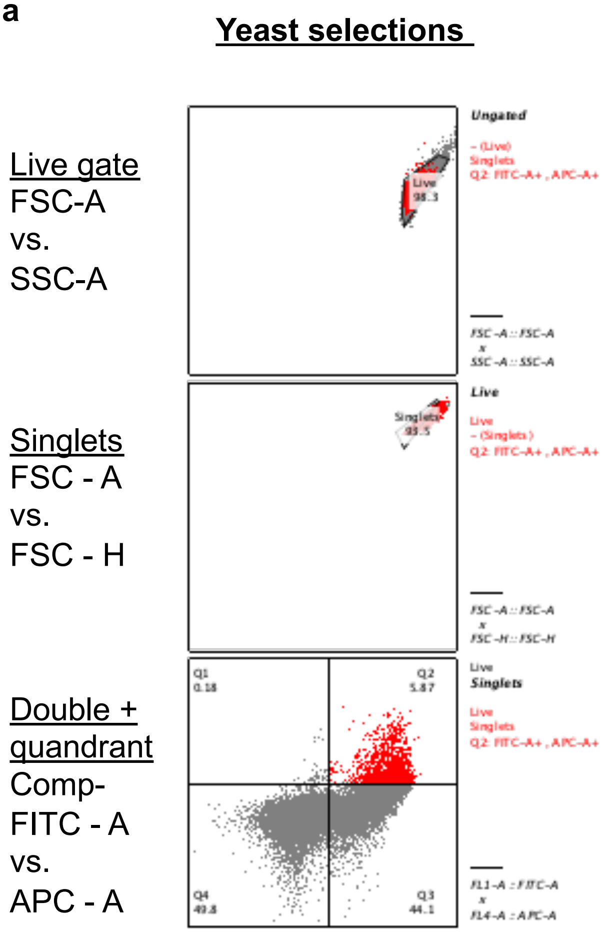


**b**

| Nanobody clone | Melting temp (℃) |
|---|---|
| A2 | 60 ± 1.4 |
| A4 | 69.7 ± 1.1 |
| A5 | 46 ± 0.23 |
| A6 | 77.2 ± 1.7 |
| B7 | 72.4 ± 0.78 |
| B8 | 77.6 ± 0.63 |
| B11 | 78.5 ± 0.46 |
| C2 | 74.8 ± 0.61 |
| C8 | 74.8 ± 0.46 |
| C9 | 69.5 ± 1.1 |
| C11 | 72.5 ± 1.3 |

Supplemental figure 2 . Quantified colony formation screen.

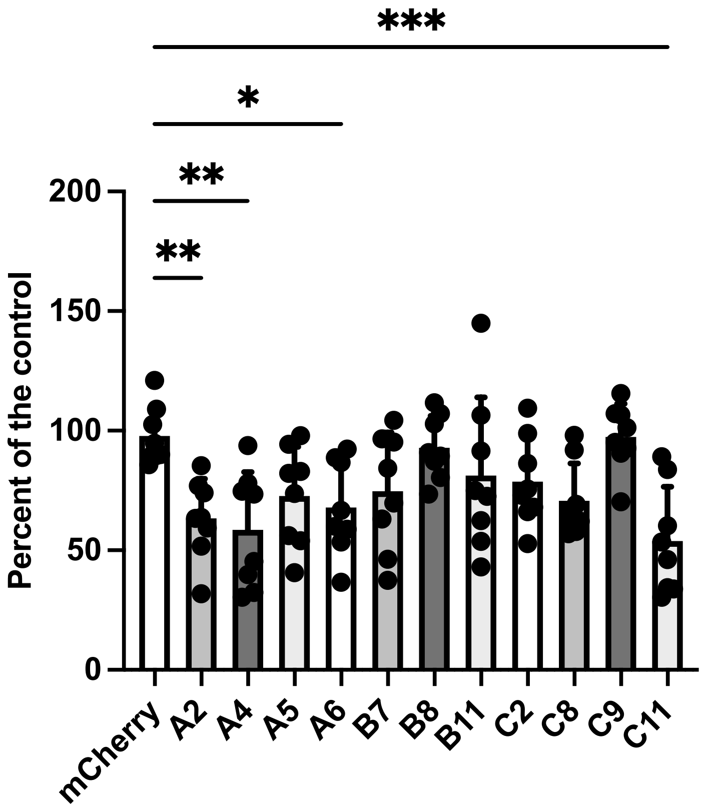

Supplemental figure 3 . Quantified Annexin-V and propidium iodide apoptosis screen.

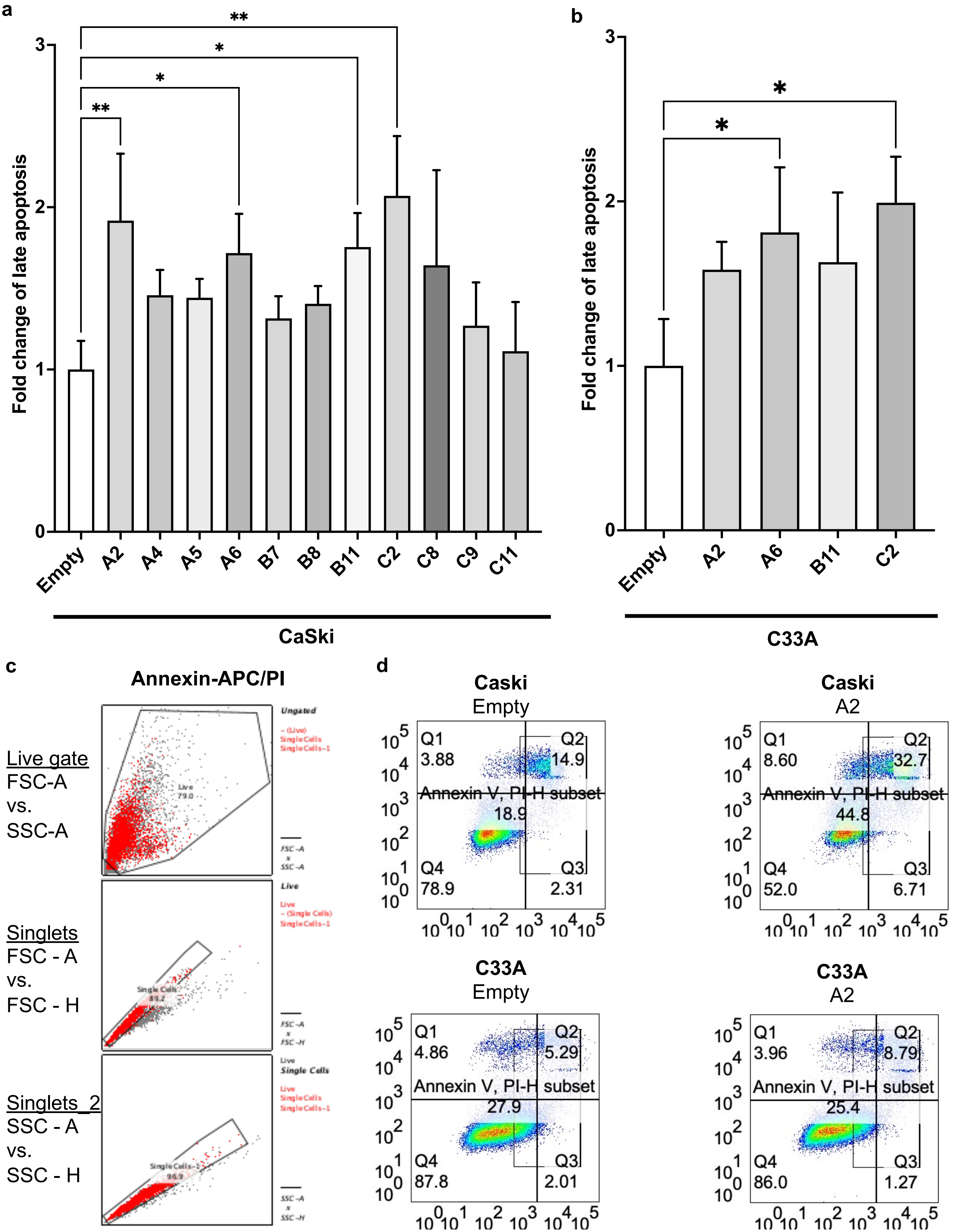

Supplemental figure 4. ISG56 screen and dose-dependent relationship of nanobody C11 on ISG56 induction.

a

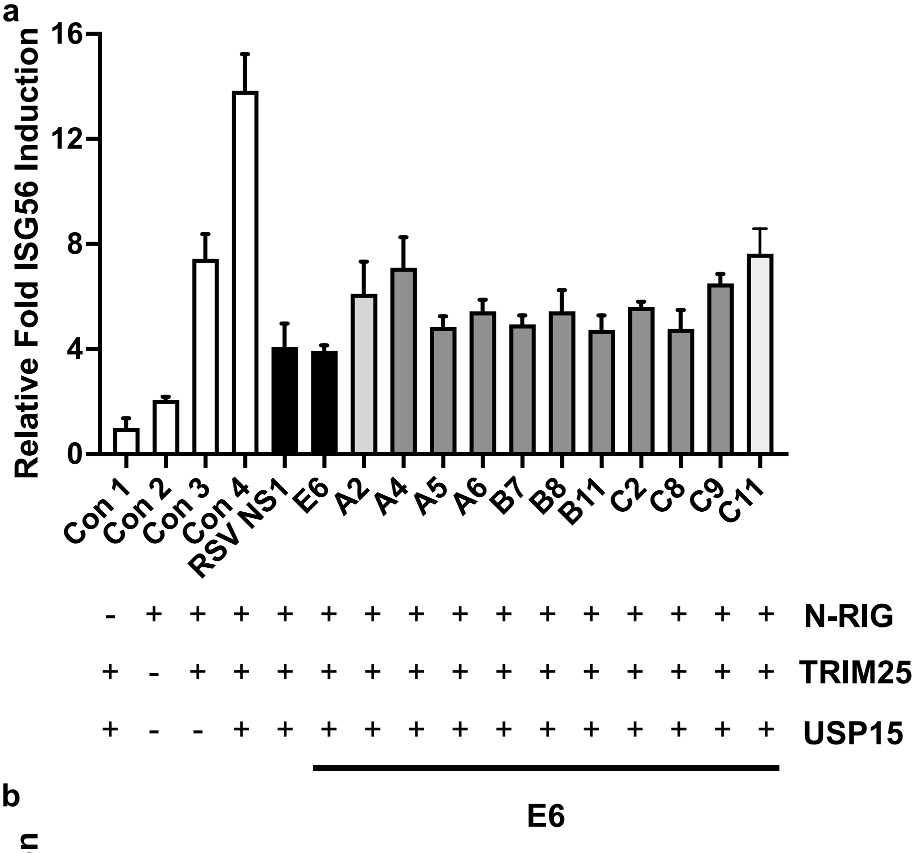


b

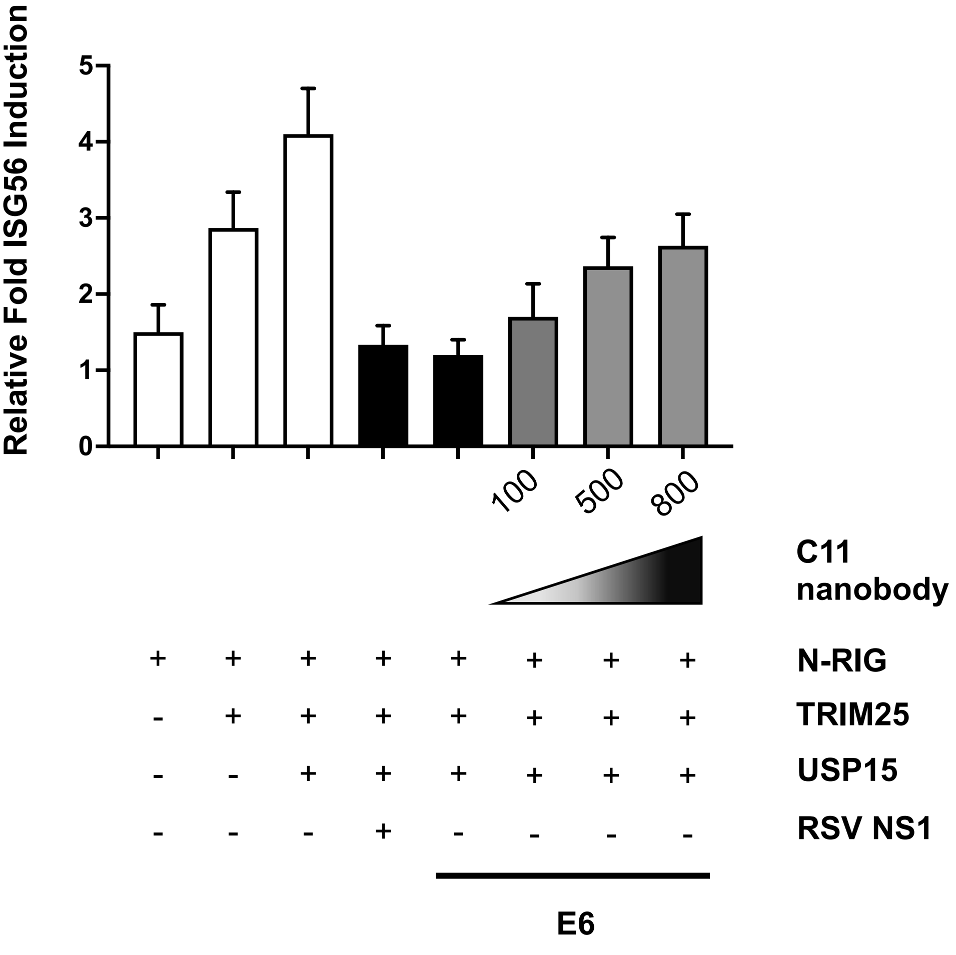

Supplemental figure 5. PROTAC$^{E6}$ induction of IFNB mRNA levels

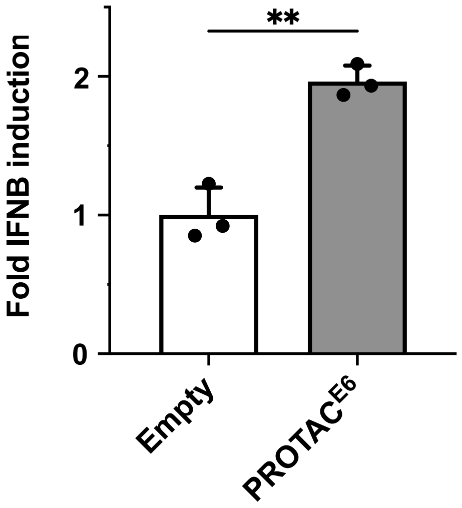

Supplemental figure 6. Mouse body weight

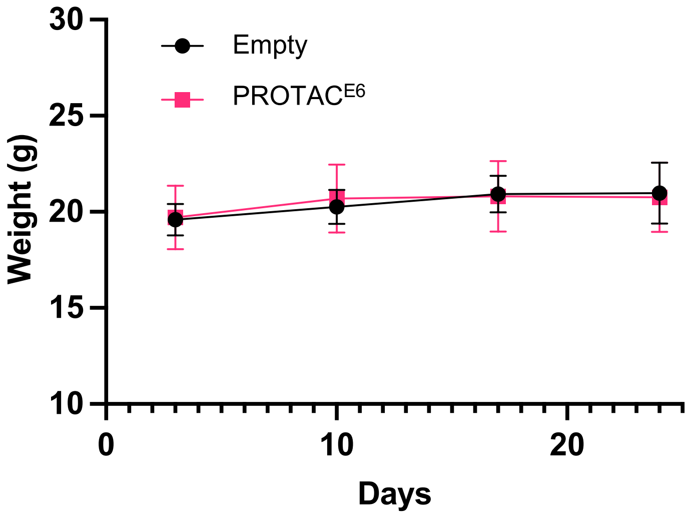
